# A chromosome-level genome assembly for the beet armyworm (*Spodoptera exigua*) using PacBio and Hi-C sequencing

**DOI:** 10.1101/2019.12.26.889121

**Authors:** Feng Zhang, Jianpeng Zhang, Yihua Yang, Yidong Wu

## Abstract

**Background:** The beet armyworm, *Spodoptera exigua* (Hübner), is a worldwide, polyphagous agricultural pest feeding on vegetable, field, and flower crops. However, the lack of genome information on this insect severely limits our understanding of its rapid adaptation and hampers the development of efficient pest management strategies.

**Findings:** We report a chromosome-level genome assembly using single-molecule real-time PacBio sequencing and Hi-C data. The final genome assembly was 446.80 Mb with a scaffold N50 of 14.36 Mb, and captured 97.9% complete arthropod Benchmarking Universal Single-Copy Orthologs (BUSCO, n=1,658). A total of 367 contigs were anchored to 32 pseudo-chromosomes, covering 96.18% (429.74 Mb) of the total genome length. We predicted 17,727 protein-coding genes, of which 81.60% were supported by transcriptome evidence and 96.47% matched UniProt protein records. We also identified 867,102 (147.97 Mb/33.12%) repetitive elements and 1,609 noncoding RNAs. Synteny inference indicated a conserved collinearity between three lepidopteran species. Gene family evolution and function enrichment analyses showed the significant expansions in families related to development, dietary, detoxification and chemosensory system, indicating these families may play a role in host plant specialization and niche adaptation.

**Conclusions:** We have generated a high-quality chromosomal-level genome that could provide a valuable resource for a better understanding and management of the beet armyworm.

## Background

*Spodoptera exigua* (NCBI:txid7107, Lepidoptera, Noctuidae), commonly known as the beet armyworm (BAW), is one of the world’s most polyphagous pests, feeding on over 90 agricultural and horticultural plants. It heavily threatens agricultural economy and food security [1–3]. The beet armyworm has been distributed throughout the tropical and temperate regions of Asia, Europe, Africa and North America although it was native to Southeast Asia [4–6]. Severe outbreaks of this pest have been recorded in some region duo to complex factors, i.e. weather (rainfall, temperature), high-level resistance and increase of host planting [5, 7]. Hence, *S. exigua* is considered an ideal model to study invasion events and population dynamics in agricultural pests [3, 4].

Currently, the use of chemical insecticides is the most widely adopted approach to suppress beet armyworm populations. However, surges in resistance to traditional insecticides (e.g. organophosphates, carbamates and pyrethroids), as well as newer insecticides with novel modes of action (e.g. emamectin benzoate, spinosad, abamectin), severely obstruct the continued use of insecticides [7, 8]. The advances in modern molecular techniques, in particular the CRISPR/Cas9 system [9, 10], have made it possible to explore the molecular functions of conserved insecticide target sites and their effect on resistance in *S. exigua* [11, 12]. However, beyond these conserved target sites, it is difficult to uncover the molecular mechanisms of resistance starting from a phenotype alone, without some knowledge of the genome context and structure. Despite being an important pest, limited genomic resources have been available until now for the beet armyworm.

Here, we present a de novo chromosome-level genome assembly of *S. exigua* from a Chinese susceptible strain using single-molecule real-time (SMRT) Pacific Bioscience (PacBio) long reads and Hi-C data. Mitochondrial genome and transcripts were also assembled. We annotated the essential genomic elements, i.e. repetitive sequences, protein-coding genes, and non-coding RNAs (ncRNAs). These valuable genomic data will help in the discovery of novel molecular mechanisms of resistance and will facilitate the design of future pest management strategies.

## Methods

### Sample collection and sequencing

The non-inbred strain WH-S, originally obtained from Wuhan Institute of Vegetables (Hubei, China) in 2009, has been maintained in the laboratory without exposure to any insecticides since its initial collection in 1998 from Wuhan [7]. Because the heterogeneity estimated from genome survey was not high (see below), WH-S strain was used for subsequent sequencing and genome assembly. Two female pupae, three adult males and six adult females were collected for sequencing experiments: one pupa for Illumina and PacBio whole genome, one pupa for Hi-C, and others for transcriptome, respectively. Genomic DNA/RNA extraction, library preparation and sequencing were carried out at BGI (Shenzhen, China) except for Hi-C library. Genomic DNA was extracted using the Qiagene Blood & CELL Culture DNA mini Kit. Long read libraries were constructed with an insert size of 15 kb using a SMRTbell DNA Template Prep Kit 1.0 and sequenced on a PacBio Sequel system (RRID:SCR_017989) using Sequel Sequencing kit v2.1 chemistry. Short read libraries of an insert size of 350 bp were prepared using the TruSeq DNA PCR-Free LT Library Preparation Kit and were subject to the paired-end 150 bp (PE 150) sequencing on the HiSeq X Ten platform (RRID:SCR_016385). Hi-C library preparation, including cross-linking the DNA, restriction enzyme digestion, end repair, DNA cyclization, and DNA purification, was performed by Frasergen Co. Ltd. (Wuhan, China). Restriction Enzyme MboI was used to digest DNA for the Hi-C assay. Genomic RNA was extracted using TRIzol™ Reagent and converted into libraries using the TruSeq RNA v2 Kit.

### Genome assembly

Quality control was performed for the raw Illumina short reads using BBTools suite v38.49 [13]: duplicates were removed using clumpy.sh; low-quality bases were trimmed using bbduk.sh, i.e. a minimum Phred quality score of 20 for both read sides, a minimum read/Ns length of 15/5 bp, a maximum length poly-A/G/C tail of 10 bp. Preliminary genome survey was investigated employing the strategy of short-read k-mer distributions using GenomeScope v1.0.0 (GenomeScope, RRID:SCR_017014) [14]. The histogram of k-mer frequencies was estimated with 21-mers using khist.sh (BBTools). Estimated genome sizes increased with the parameter of maximum k-mer coverage cutoff, thus we selected 1,000 and 10,000 as the maximum k-mer coverage cutoffs.

The raw PacBio long reads were assembled into contigs using Flye v2.5 (Flye, RRID:SCR_017016) [15] and Falcon v1.3.0 (Falcon, RRID:SCR_016089) [16]; both assemblies were self-polished with a round of Flye polishing module “--polish-target”. To improve contig contiguity, Flye and Falcon assemblies were merged with two rounds of quickmerge v.0.3 [17] following USAGE 2 [18]; Falcon assembly was treated as the query in both rounds because of its higher contiguity (greater N50 value); Flye assembly was treated as the reference in the first round and the merged assembly from the first round was set as the reference in the second round; for each round, the N50 length of the reference assembly was assigned to the length cutoff for anchor contigs (‘-l’). We removed redundant heterozygous contigs with Purge Haplotigs v1.1.0 [19]; percent cutoff for identifying a contig as a haplotig was set as 50 (-a 50) with other parameters as the default. Non-redundant assembly was further polished with Illumina short reads using two rounds of Pilon v1.22 (Pilon, RRID:SCR_014731 [20]; input BAM mapping files were generated using Minimap2 v2.17 (Minimap2, RRID:SCR_018550) [21] and SAMtools v1.9 (Samtools, RRID:SCR_002105) [22].

To perform the chromosome-level concatenation, we aligned Hi-C read pairs to the genome, removed duplicates, and extracted the Hi-C contacts using Juicer v1.6.2 (Juicer, RRID:SCR_017226) [23]. Pseudo-chromosomes were generated using 3D-DNA v180922 [24] in combination with a review step in Juicebox Assembly Tools module within Juicebox [23]. Sex chromosomes with female heterogamety were determined according to male-to-female comparison approach [25]. In theory, the average depth of sex chromosomes should be about half of that for autosomes in females, and the average depth of Z chromosome should be approximately equal to autosomes whereas W chromosome should nearly be zero in males. Firstly, we mapped Illumina short reads to the genome assembly using Minimap2 [21], then the average depth of coverage was calculated using SAMtools [22]. In fact, the average depth of coverage of W chromosome is higher than expected because harboring genes may be similar to some regions of autosomes and Z chromosomes [25]. Chromosomal nomenclature order and orientation were assigned in comparison with *Bombyx mori* genome [26].

Contaminants were removed against the nt and UniVec databases using HS-BLASTN v0.0.5 [27] and BLAST+ (i.e. blastn) v2.7.1 (BLASTN, RRID:SCR_001598) [28]. We assessed the assembly completeness using Benchmarking Universal Single-Copy Orthologs (BUSCO, RRID:SCR_015008) [29] analyses against insect reference dataset (insect_odb9, n = 1,658). We also mapped PacBio long reads and Illumina short reads to the final genome assembly with Minimap2.

### Genome annotation

Repetitive elements, ncRNAs and protein-coding genes were annotated for the beat armyworm. To identify repeat elements, we constructed a de novo species specific repeat library using RepeatModeler v1.0.11 (RepeatModeler, RRID:SCR_015027) [30], and then combined it with Dfam_3.0 [31] and RepBase-20181026 databases [32] to generate a custom library. We then used RepeatMasker v4.0.9 (RepeatMasker, RRID:SCR_012954) [33] to mask repeats in the genome assembly with the combined custom library. Non-coding RNAs were identified with Infernal v1.1.2 (Infernal, RRID:SCR_011809) [34] and tRNAscan-SE v2.0 (tRNAscan-SE, RRID:SCR_010835) [35].

Protein-coding genes were predicted with the MAKER v2.31.10 pipeline (MAKER, RRID:SCR_005309) [36] which can integrate ab initio, transcriptome- and protein homology-based evidence. For the ab initio gene predictions, we trained gene model parameters for Augustus v3.3 (Augustus, RRID:SCR_008417) [37] and GeneMark-ET v4.38 (GeneMark, RRID:SCR_011930) [38] using BRAKER v2.1.0 [39], which incorporated RNA-seq data to accurately identify exon/intron boundaries. Input BAM alignment file for BRAKER was generated with HISAT2 v2.1.0 (HISAT2, RRID:SCR_015530) [40]. For the transcriptome-based evidence, a genome-guided method was applied to transcript assembly using StringTie v1.3.4 (StringTie, RRID:SCR_016323) [41]. For the protein homology-based evidence, protein sequences of *Apis mellifera* (GCF_003254395.2), *Bombyx mori* (GCF_000151625.1), *Drosophila melanogaster* (GCF_000001215.4), *Tribolium castaneum* (GCF_000002335.3), *Helicoverpa armigera* (GCF_002156985.1) and *Spodoptera litura* (GCF_002706865.1) were downloaded from the National Center for Biotechnology Information (NCBI) [42]. Finally, repeat-masked genome and repeat annotations, assembled transcripts, protein sequences, and ab initio predicted gene models from BRAKER were passed to MAKER for the gene structure predictions.

Gene functions were assigned to MAKER predicted gene models using Diamond v0.9.24 (DIAMOND, RRID:SCR_016071) [43] against the UniProtKB (UniProtKB, RRID:SCR_004426) database [44] with the sensitive mode ‘--more-sensitive −e 1e-5’. We searched protein domains, Gene Ontology (GO) and Reactome pathway using InterProScan 5.41-78.0 (InterProScan, RRID:SCR_005829) [45] against Pfam (Pfam, RRID:SCR_004726) [46], Gene3D (Gene3D, RRID:SCR_007672) [47], Superfamily (SUPERFAMILY, RRID:SCR_007952) [48], CDD (Conserved Domain Database, RRID:SCR_002077) [49], and SMART (SMART, RRID:SCR_005026) [50] databases (‘-dp −f TSV, GFF3 -goterms -iprlookup - pa -t p -appl Pfam, Smart, Gene3D, Superfamily, CDD’). We also annotated genes using eggNOG-mapper v2.0 [51] against the eggNOG v5.0 (eggNOG, RRID:SCR_002456) database [52] to obtain additional information of GO, Kyoto Encyclopedia of Genes and Genomes (KEGG, RRID:SCR_012773) orthology (ko), KEGG pathways, enzyme codes (ECs), and clusters of orthologous groups (COG, RRID:SCR_007139). Circus figure showing the element tracks was produced by Tbtools v1.0 [53].

### Gene family evolution

We selected 14 species for orthogroup inference, including Diptera (*Drosophila melanogaster*), Trichoptera (*Stenopsyche tienmushanensis*), Plutellidae (*Plutella xylostella*), Pyralidae (*Amyelois transitella*), Nymphalidae (*Danaus plexippus*), Geometridae (*Operophtera brumata*), Sphingidae (*Manduca sexta*), Bombycidae (*Bombyx mori*), Noctuidae (*Helicoverpa armigera*, *Heliothis virescens*, *Spodoptera exigua*, *Spodoptera frugiperda*, *Spodoptera litura*, *Trichoplusia ni*). Lepidoptera taxa covered a butterfly and six representative pest families. All protein sequences were downloaded from the NCBI except for *Stenopsyche tienmushanensis* from GigaScience Database [54]. Orthogroups were inferred using OrthoFinder v2.3.8 (OrthoFinder, RRID:SCR_017118) [55] with the Diamond as the sequence aligner.

Phylogenomic reconstructions were performed using single-copy orthologous inferred from OrthoFinder. Protein sequences of each orthogroup were aligned using MAFFT v7.394 (MAFFT, RRID:SCR_011811) [56] with the L-INS-I strategy, trimmed using trimAl v1.4.1 (trimAl, RRID:SCR_017334) [57] with the heuristic method ‘automated1’, and concatenated using FASconCAT-G v1.04 [58]. Phylogenetic tree was reconstructed using IQ-TREE v2.0-rc1 (IQ-TREE, RRID:SCR_017254) [59] with the substitution model restricted to the LG model and 10% of partition pairs for partition clustering algorithm to reduce the computational burden (‘-m MFP --mset LG --msub nuclear --rclusterf 10’). Genes that violate models, i.e. genes undergoing non-SRH (stationary, reversible, and homogeneous) evolution, were also removed prior to tree inference (‘--symtest-remove-bad --symtest-pval 0.10’). Node support values were calculated (‘-B 1000 --alrt 1000’) using 1,000 SH-aLRT replicates [60] and 1,000 ultrafast bootstraps [61]. Divergence time was inferred using MCMCTree within the PAML v4.9j (PAML, RRID:SCR_014932) package [62]. Four fossil calibrations were extracted from the Paleobiology Database [63] for dating: root (314.6‒318.1 Mya), Trichoptera (311.4‒314.6 Mya), Ditrysia (93.5‒145.2 Mya) and Noctuoidea (28.1‒33.9 Mya).

Expansions and contractions of gene families were estimated using CAFÉ v4.2.1 [64] using the method of single birth-death parameter lambda with the significance level of 0.01. Function enrichment analyses of GO and KEGG categories were also performed for those significantly expanded families using R package clusterProfiler v3.10.1 (clusterProfiler, RRID:SCR_016884) [65] with the default significance values (p-value as 0.01 and q-value as 0.05).

### Genome synteny

To search for syntenic blocks between genomes, coding sequences (CDSs) of *S. exigua* (except for the W chromosome) and other two species (*B. mori*, *S. litura*) were used for analyses. Chromosome W was not included because *S. litura* genome did not assign any contigs into this chromosome. Chromosome-level genomic resources were downloaded from the NCBI (*S. litura*, GCF_002706865.1) and SilkBase v2.1 (*B. mori*) [66]. We searched pairwise synteny using LAST v1080 (LAST, RRID:SCR_006119) [67] and removed tandem duplications and weak hits by the module McScan (MCScan, RRID:SCR_017650) [68] (Python version) in jcvi [69]. We extracted subset of blocks with following options “--minspan=30 -- minsize=15 --simple”. Finally, the high-quality syntenic blocks were visualized by Circos v0.69-9 (Circos, RRID:SCR_011798) [70].

## Results

### Genome assembly

We generated 3,919,515 PacBio subreads of 38.98 Gb (87X) with the mean and N50 length reaching 9.95 and 16.01 kb, respectively. A total of 34.62 Gb (77X), 65.21 Gb and 56.32 Gb (126X) were produced in the Illumina X Ten platform for whole genome, transcriptome and Hi-C sequencing, respectively. Estimated genome size ranged from 408.58 Mb to 448.90 Mb (Table S1). Genome repeat length increased with the maximum k-mer coverage cutoff, ranging from 64.96 Mb to 104.93 Mb. Heterozygosity rate (0.585‒0.594) and a small indistinct peak on the left of main peak (Fig. 1a) implied that the heterozygosity of the sequenced strain was not high.

**Figure 1:**
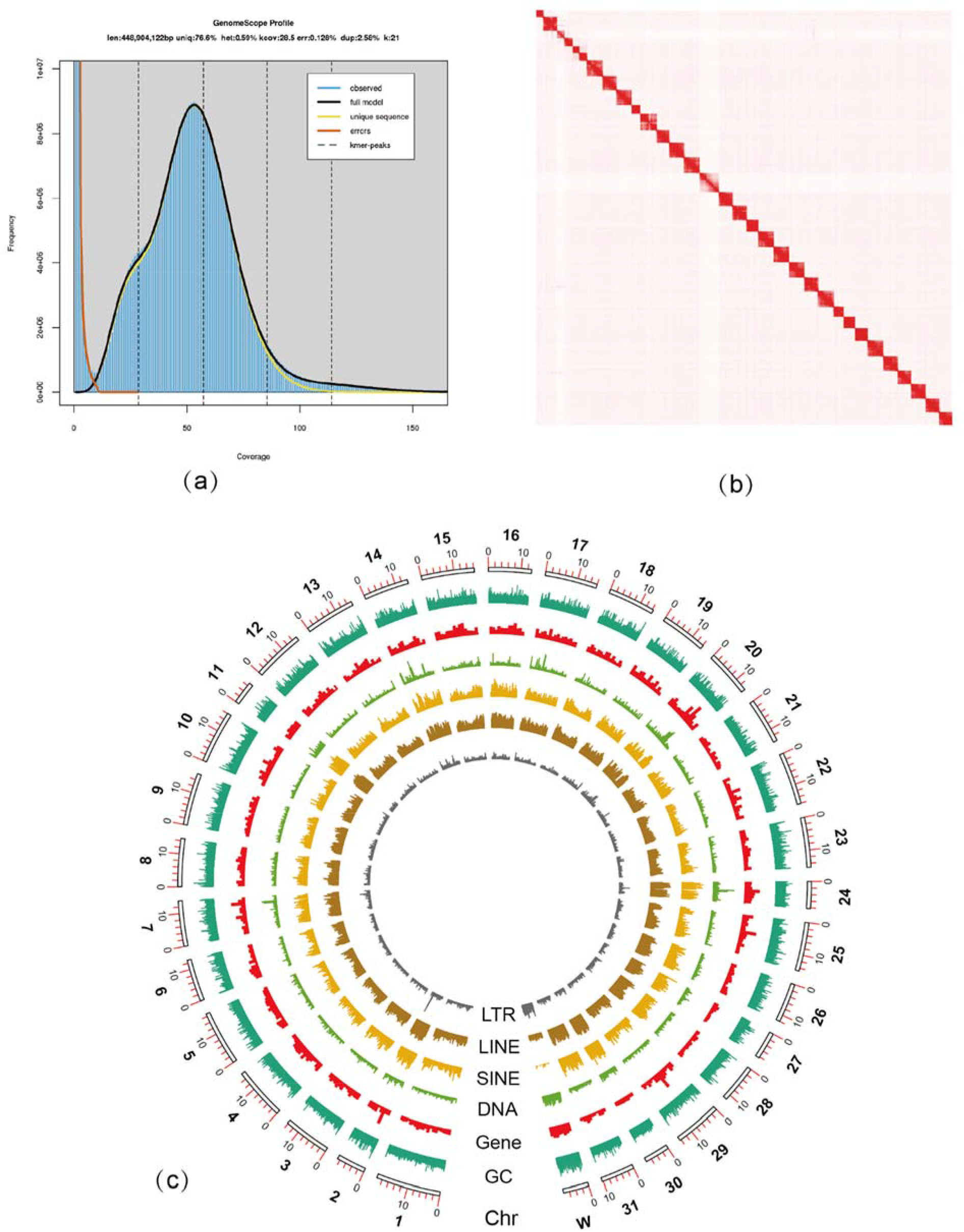
Genome characteristics of the beet armyworm genome. (a) Genome survey from 21-k-mer frequency distribution plot estimated using GenomeScope (a maximum k-mer coverage of 10,000). (b) Hi-C contact map with and among 32 curated pseudo-chromosomes (squares) generated from Juicebox. (c) Circos tracks showing element distribution from outside to inside: GC content, gene density, DNA transposons, SINEs, LINEs and LTRs; 10 kb sliding windows for GC content and 100 kb for others; the most outer numbers indicating the chromosome nomenclature order and sizes (Mb).

Both Flye and Falcon assemblies (Table S2) were ca. 100 Mb larger than the estimated size due to a low-moderate rate of heterozygosity (duplicated BUSCOs > 10%), and merged into a 561.68 Mb-assembly of 934 contigs (N50 length 3.47 Mb). We removed 668 redundant heterozygous sequences using Purge Haplotigs, resulting an obvious improvement of N50 length (5.53 Mb). After polishing, chromosome concatenation and contaminant removal, the final genome assembly had 301 scaffolds/667 contigs, a total length of 446.80 Mb, scaffold/contig N50 length of 14.36/3.47 Mb, and a GC content of 36.67%; 367 contigs were concatenated into 32 pseudo-chromosomes (Fig. 1b), accounting for 96.18% (429.74 Mb) of the total genome length. Through 3D-DNA pipeline, partial assembled scaffolds or contigs in previous assembly process were split into shorter sequences due to the misjoints. We successfully determined 30 autosomes, Z and W chromosome with their nomenclature order and orientation homologous to those of *B. mori* (Fig. 1c). The final assembled size was very close to the estimated size from GenomeScope (Table S1). BUSCO completeness analyses against insect_odb9 reference dataset (n=1,658) (Table S2) showed our assembled versions covered 97.8‒98.6% complete, 0.2‒0.4% fragmented, and 1.1‒1.9% missing BUSCO genes. We mapped 97.62% and 93.78% Illumina short and PacBio long reads to our final assembly. All these assessments showed the accuracy and the high integrity of our final genome assembly with low redundancy (2.1 % duplicated BUSCOs). Among four Noctuidae pest species (Table 1), assemblies of *S. exigua* and *S. frugiperda* have the highest genome contiguity (contig N50 > 1 Mb, contig number < 1,000) and the lowest gap content (< 0.1%).

**TABLE 1:** Genome assembly statistics of four Noctuidae species available on GenBank

| Characteristics | *Spodoptera*<br>*exigua* | *Spodoptera*<br>*litura* | *Spodoptera*<br>*frugiperda* | *Helicoverpa*<br>*armigera* |
| --- | --- | --- | --- | --- |
| Accession number | GCA_011316535.1 | GCA_002706865.1 | GCA_011064685.1 | GCA_002156985.1 |
| Assembly size (Mb) | 446.80 | 438.96 | 486.29 | 337.09 |
| Number of scaffolds | 301 | 2,975 | 92 | 998 |
| Longest scaffold (Mb) | 19.74 | 18.94 | 22.37 | 6.15 |
| Scaffold N50 length (Mb) | 14.36 | 13.59 | 16.35 | 1.00 |
| Number of contigs | 667 | 13,637 | 617 | 24,552 |
| Longest contig (Mb) | 15.88 | 0.64 | 16.14 | 0.31 |
| Contig N50 length (Mb) | 3.47 | 0.068 | 1.13 | 0.023 |
| Pseudo-chromosomes | 32 | 31 | 32 | 0 |
| GC (%) | 36.67 | 36.56 | 36.43 | 37.50 |
| Gaps (%) | 0.075 | 2.49 | 0.011 | 11.01 |
| BUSCO completeness (%) | 97.9 | 97.1 | 93.4 | 96.8 |

### Genome annotation

By combing *de novo* and homology-based evidence, we identified that 33.12% (147.97 Mb) of the genome was composed of repetitive elements in the beet armyworm genome. Repeat ratio was very close to the repeat content (31.8%) in the published *S. litura* genome (Cheng et al. 2017). The main interspersed repeat families were LINE (14.81%), Unknown (5.89%), RC (3.95%), LTR (3.10%), SINE (2.53%), and DNA (1.79%). Simple repeats also occupied 0.83% of the genome (Table S3). Interestingly, chromosome W had more abundant DNA transposons and LTR retrotransposons but fewer interspersed nuclear elements (SINEs and LINEs) compared to other chromosomes (Fig. 1c).

We identified 172 rRNAs (5S, 5.8S, LSU, SSU), 61 miRNAs (37 families), 118 small nuclear RNAs (snRNAs, 19 families), 2 long non-coding RNAs (lncRNA-Sphinx), 1,135 transfer RNAs (tRNAs), 3 ribozymes (2 families), 114 cis-regulatory elements, and 3 other (Metazoa_SRP) ncRNAs (Table S4). Twenty-one tRNA isotypes were identified except for SelCys-isotype which was lacking. Small nuclear RNAs were classified into six spliceosomal RNAs (U1, U2, U4, U5, U6, U11), three minor spliceosomal RNAs (U12, U4atac, U6atac), and four H/ACA box and 20 C/D box small nucleolar RNAs (snoRNAs).

We predicted 17,727 protein-coding genes (gene models) with a mean length of 9,604.99 bp in the beet armyworm genome. BUSCO completeness assessment using the protein mode (‘-m prot’) identified 1,601 (96.6%) complete, 45 (2.7%) duplicated, 12 (0.7%) fragmented, and 45 (2.7%) missing BUSCO genes, indicating the highly completeness of our predicted protein-coding gene set. Exons and introns had a mean count of 6.40 and 5.20 per gene and a mean length of 334.78 bp and 1,434.55 bp, respectively (Table 2). Among predicted genes, 81.60% were supported by transcriptome evidence and 96.47% matched the UniProt protein records. InterProScan identified protein domains for 13,186 (74.38%) genes, GO terms for 8,227 (46.69%) genes and Reactome pathways for 2,731 (15.41%) genes, respectively. eggNOG identified GO, KEGG orthology (ko), KEGG pathway, ECs, and COG for 6,706, 7,643, 4,493, 2,675 and 12,987 genes, respectively.

**TABLE 2:** Genome annotation statistics of the beet armyworm

| Elements | Number |
| --- | --- |
| Protein-coding genes | 17,727 |
| Mean protein (aa) and gene (bp) length | 482.19/9,604.99 |
| Exons/introns per gene | 6.40/5.20 |
| Exon | 37.95 Mb (8.49%) |
| Intron | 132.31 Mb (28.34%) |
| Mean exon/intron length | 334.78/1,434.55 |
| BUSCO completeness (%) | 96.6 |
| GO | 10,298 |
| Reactome pathway | 2,731 |
| KEGG ko | 7,643 |
| KEGG pathway | 4,493 |
| EC | 2,675 |
| COG | 12,987 |

### Gene family evolution

We assigned a total of 95.59% (214,213) genes into 15,906 orthogroups (gene families) using OrthoFinder. Among them, 4,618 orthogroups were shared by all fourteen species and 1,215 are single-copy ones; 62 families and 586 orthologs are common to six Noctuidae species; 2,376 orthogroups and 9,324 orthologs were species-specific. For the beat armyworm, 17,094 (96.43%) genes were assigned into 11,663 orthogroups; 154 families and 569 genes were species-specific (Fig. 2a, Table S5).

**Figure 2:**
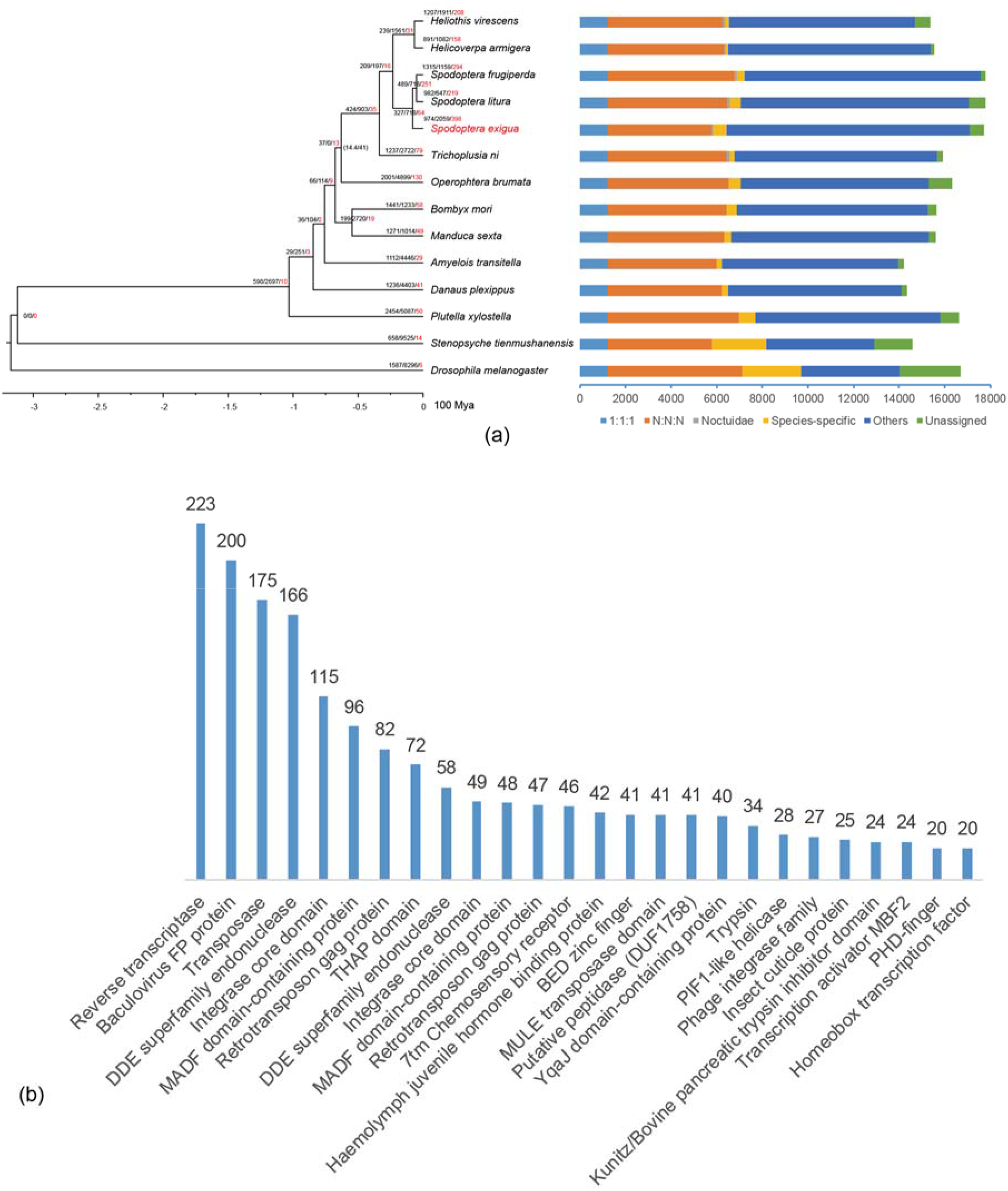
Phylogeny and gene family evolution. (a) Dating tree with node values representing the number of expanded, contracted and rapidly evolving families. Nodes support values fewer than 100 indicated within brackets. ‘1:1:1’ as shared single-copy genes, ‘N:N:N’ as orthologs from shared multi-copy orthogroups, ‘Noctuidae’ as shared orthologs unique to Noctuidae, ‘Others’ as unclassified orthologs, ‘Unassigned’ as orthologs which cannot be assigned into any orthogroups. (b) The significantly expanded families with gene number of the family greater than 20.

For phylogenomic analyses, we removed 109 single-copy loci that violated SRH hypothesis, resulting a matrix of 1,106 loci and 456,865 amino acid sites. Our phylogeny was largely consistent with the newest lepidopteran classification [71] except for the position of *Operophtera brumata* (Geometridae), which was clustered to Noctuoidea than Bombycoidea. Very weak node support (14.4/41) also confirmed the unstable position of this species (Fig. 2a). Divergence time estimation showed that Noctuidae diverged 32.58~34.47 Mya, conforming to the oldest Noctuidae fossil (*Noctuites incertissima*, 28.1~33.9 Mya) [72]. The bear armyworm diverged from the other two *Spodoptera* species 7.40~8.47 Mya.

We identified 974/2,059 expanded/contracted gene families in *S. exigua* genome (Fig. 2a), 278 and 120 of them being rapidly evolving at the significant level of 0.01. Haemolymph juvenile hormone binding protein (JHBP), chorion protein and insect cuticle protein have a profound effect in regulating embryogenesis, maintaining the status quo of larva development and stimulating adult reproductive maturation [73–75]. Gustatory receptors (GRs, referred as 7tm chemosensory receptor in Fig. 2b) were also found to be expanded in sampled 14 species (Fig. 2b). Massive expansion of GRs in Lepidoptera may be related to extreme polyphagy [76, 77]. *S. exigua* feeds a large number of host plants including more than 40 monocot families [78]. Trypsin and cytochrome P450 (CYP, not shown in Fig. 2b) plays an important role in digestion [79] and detoxification [80] by contributing to protein digestion, xenobiotic metabolism, insecticide resistance, odorant or pheromone metabolism. In addition, many gene families are transposon repeat-related, such as reverse transcriptase, DDE superfamily endonuclease, Baculovirus FP protein, transposase, integrase core domain-containing protein, retrotransposon gag protein, phage integrase etc. Large expansions of these transposable elements explain the relatively high ratio of repetitive elements, particularly LINE superfamilies CR1-Zenon, L2, R1 and RTE (Table S3), although their functions are largely unknown.

GO and KEGG function of rapidly expanded families furthermore refined above hypotheses. The top six GO enrichment categories, i.e. GO term count greater than 25, related to transposon and egg chorion (Fig. 3a). KEGG pathways (Fig. 3b) were enriched in development (folate biosynthesis), digestion (cholesterol metabolism, lysosome, arachidonic acid metabolism, carbohydrate digestion and absorption), and detoxification (xenobiotic and drug metabolism by CYP, glutathione metabolism). Gene family evolution provides valuable information supporting genetic mechanisms of polyphagy and ecological adaptions for the beet armyworm.

**Figure 3:**
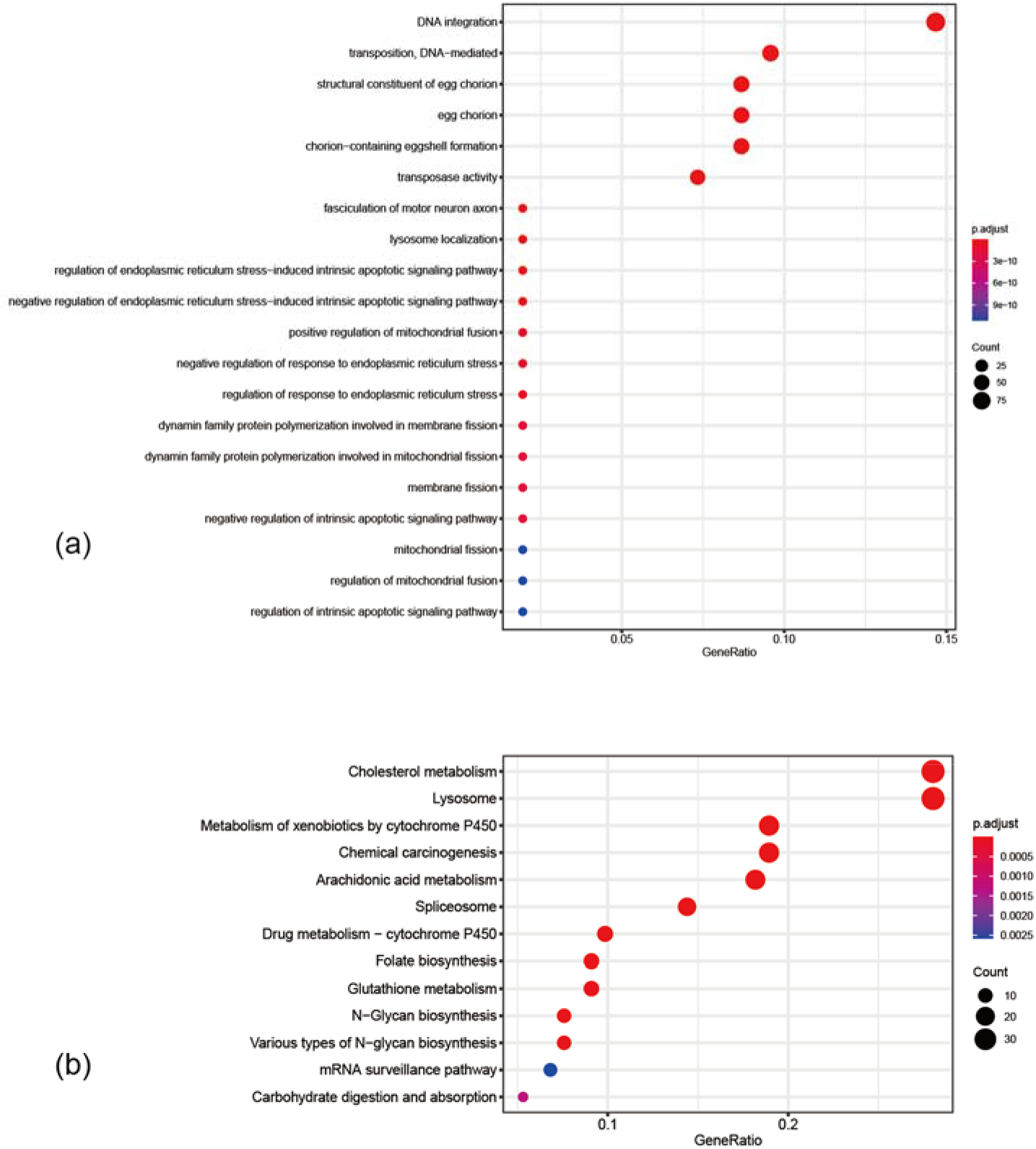
GO (a) and KEGG (b) function enrichment of significantly expanded gene families. Only the top 20 categories shown.

### Synteny

Syntenic analyses showed that 100 syntenic blocks between *S. exigua* and *B. mori* and 148 syntenic blocks between *S. exigua* and *S. litura* were conserved. Three *B. mori* chromosomes 11, 23 and 24 were clearly divided into three pairs in *S. exigua*: 11 and 29, 23 and 30, 24 and 31, respectively (Fig. 4a). Strong syntenic correspondence between *S. exigua* and *S. litura* (Fig. 4b) indicated the conserved chromosome-level gene collinearity, which was also reported previously [81, 82].

**Figure 4:**
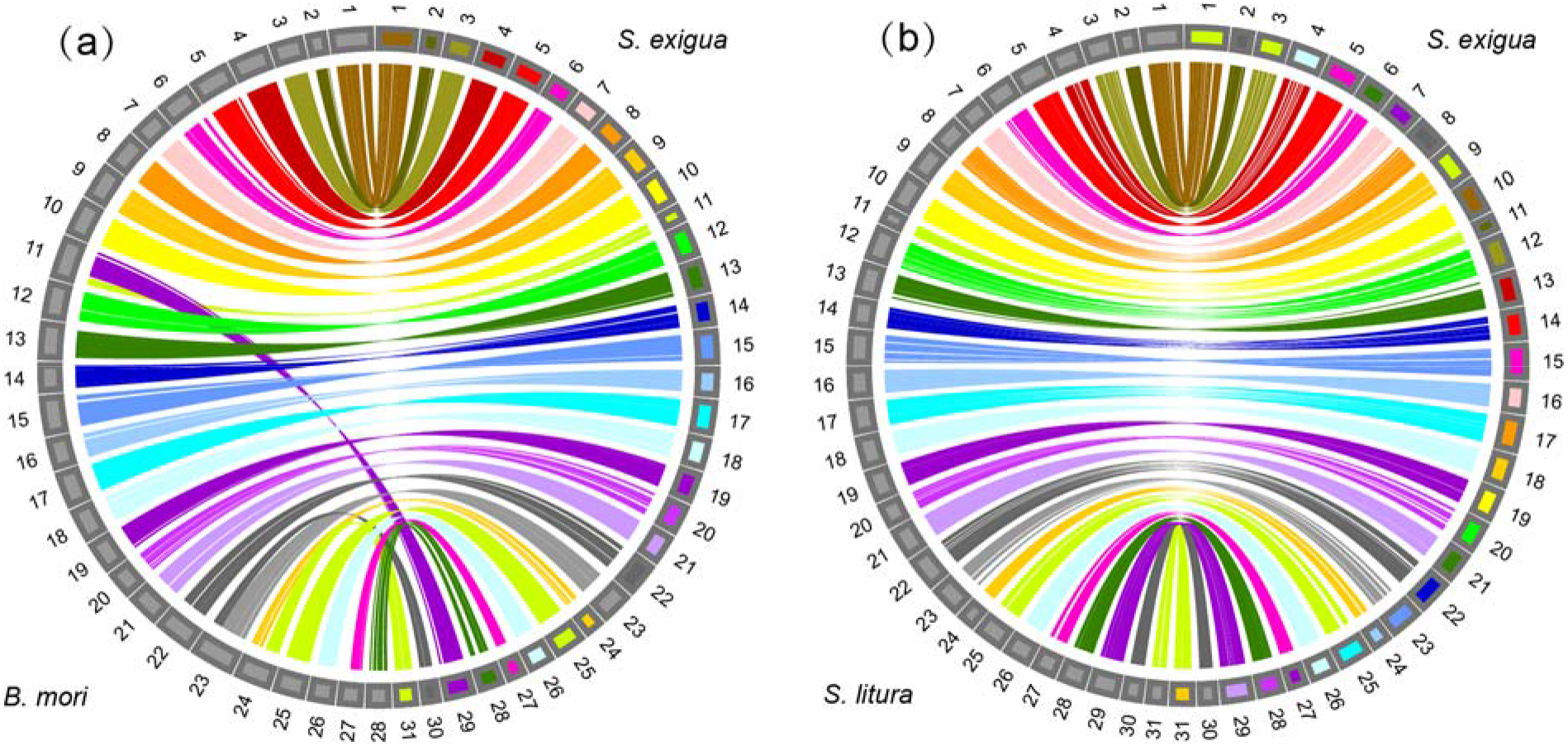
Circos plot of the synteny between the genomes. (a) *S. exigua* and *B. mori*. (b) *S. exigua* and *S. litura*.

## Availability of Supporting Data and Materials

The project and raw sequencing data have been deposited at the NCBI under the accessions PRJNA588360 and SRR10441878–SRR10441889. The data sets supporting the results of this article are available in the GigaScience GigaDB repository.

## Additional Files

**Supplementary Table S1:** Genome survey for *Spodoptera exigua*.

**Supplementary Table S2:** Summary of each assembly version for *Spodoptera exigua*.

**Supplementary Table S3:** Repeat annotation in *Spodoptera exigua*.

**Supplementary Table S4:** Summary statistics for non-coding RNAs in *Spodoptera exigua*.

**Supplementary Table S5:** Summary statistics for gene family inference.

## Abbreviations

BAM: binary alignment map
BLAST: Basic Local Alignment Search Tool
bp: base pairs
BUSCO: benchmarking universal single-copy orthologs
CDS: coding sequence
COG: clusters of orthologous group
CYP: cytochrome P450
EC: enzyme codes
Gb: gigabase pairs
GC: guanine-cytosine
GO: gene ontology
GR: gustatory receptor
Hi-C: chromosome conformation capture
kb: kilobase pairs
KEGG: Kyoto Encyclopedia of Genes and Genomes
ko: KEGG orthology
LINE: long interspersed nuclear element
lncRNA: long non-coding RNA
LSU: large subunit rRNA
LTR: long terminal repeat
Mb: megabase pairs
miRNA: micro RNA
ncRNA: non-coding RNA
NCBI: National Center for Biotechnology Information
PacBio: Pacific BioSciences
PBDB: Paleobiology Database
PE: paired end
RC: rolling circle
RNA-seq: RNA sequencing
rRNA: ribosomal RNA
SINE: short interspersed nuclear element
SMRT: single-molecule real time
snRNA: small nuclear RNA
snoRNA: small nucleolar RNA
SRA: Sequence Read Archive
SSU: small subunit rRNA
tRNA: transfer RNA

## Competing Interests

All authors declare that they have no competing interests.

## Funding

This work was supported by a grant (no. 31572030) from the National Natural Science Foundation of China and the Fundamental Research Funds for the Central Universities (KYZ201920).

## Authors’ Contributions

Y.W. and Y.Y. conceived this study. F.Z. and J.Z. carried out the experiments and performed the analyses. F.Z. and Y.W. took the lead in writing the manuscript. All authors discussed the results and contributed to the final manuscript.

## Notes

### Competing Interest Statement

The authors have declared no competing interest.

